# EEG frequency tagging reveals neural entrainment to people moving in synchrony

**DOI:** 10.1101/2020.11.06.368118

**Authors:** Emiel Cracco, Haeeun Lee, Goedele van Belle, Lisa Quenon, Patrick Haggard, Bruno Rossion, Guido Orgs

## Abstract

Humans and other animals have evolved to act in groups, but how does the brain distinguish multiple people moving in group from multiple people moving independently? Across three experiments, we test whether biological motion perception depends on the spatiotemporal relationships among people moving together. In Experiment 1, we apply EEG frequency tagging to apparent biological motion and show that fluently ordered sequences of body postures drive brain activity at three hierarchical levels of biological motion processing: image, body sequence, and movement. We then show that movement-, but not body- or image-related brain responses are enhanced when observing four agents moving in synchrony. Neural entrainment was strongest for fluently moving synchronous groups (Experiment 2), displayed in upright orientation (Experiment 3). Our findings show that the brain preferentially entrains to the collective movement of human agents, deploying perceptual organization principles of synchrony and common fate for the purpose of social perception.

## Introduction

Both human (Dyer et al., 2009; Raafat et al., 2009) and non-human animals (Couzin, 2018; Sumpter, 2006) have evolved to move in groups. In birds, flying in V-formation does not just provide safety in numbers, it also saves energy (Weimerskirch et al., 2001). In humans, performing dance or music together fosters cooperation, prosocial behavior, and group affiliation (Grau, 2015; Rennung & Göritz, 2016; Tarr et al., 2015; von Zimmermann et al., 2018). The same is true for perceiving coordinated behavior: Watching other people move in sync activates reward-related brain areas (Eskenazi et al., 2015), signals affiliation (Lakens & Stel, 2011; Marques-Quinteiro et al., 2019; Wilson & Gos, 2019), and adds to the aesthetic appreciation of both music (Hagen & Bryant, 2003) and dance (Vicary et al., 2017).

However, while the adaptive functions of group behavior are well-documented, little is known about how the brain represents the movements of groups (Riddell & Lappe, 2018; Sweeny et al., 2013). Indeed, existing models of biological motion perception have largely focused on processing the movements of individuals (Giese & Poggio, 2003) or pairs (Hovaidi-Ardestani et al., 2018). These models have distinguished two pathways for processing biological movement. In the *structure-from-motion* pathway, biological motion perception arises from the kinematics of the observed actions; in the *motion-from-structure* pathway, it arises from combining sequences of static body shape snapshots into coherent movement (Giese & Poggio, 2003; Lange & Lappe, 2006). This dual pathway structure is supported by evidence that biological motion perception does not require moving stimuli but can also be induced by sequences of static body images (Orgs et al., 2011, 2013; Orgs & Haggard, 2011), through neurons that integrate static body information over time (Jellema & Perrett, 2003; Singer & Sheinberg, 2010), both in extra-striate body- and movement specific brain areas and in motor areas of the brain (Downing et al., 2006; Orgs et al., 2016; Stevens et al., 2006).

Thus, research indicates that processing individual actions involves temporal integration of static body snapshots into movements (Giese & Poggio, 2003; Lange & Lappe, 2006). But how do these mechanisms contribute to group perception? Speaking to this question, recent studies have shown that the actions of multiple people can be processed in parallel (Cracco et al., 2015, 2016, 2019; Cracco & Brass, 2018a, 2018b, 2018c; Cracco & Cooper, 2019; Tsai et al., 2011). However, whether multiple people form a group is determined not by the number of people but primarily by the relationships among their movements (Templeton et al., 2018). Hence, a key question is how the brain distinguishes multiple people moving together from multiple people moving alone. Here, we test the hypothesis that this involves two hierarchical stages of biological motion perception. First, body postures are combined into movements (Giese & Poggio, 2003; Lange & Lappe, 2006). Next, these movements are bound into groups, by employing perceptual grouping principles such as synchrony and common fate (Wagemans et al., 2012) for the purpose of social perception.

To test this hypothesis, we developed a new EEG paradigm that dissociates different levels of biological motion processing. Specifically, we combined apparent biological motion (Orgs et al., 2011, 2013) with frequency tagging (Norcia et al., 2015) by letting participants watch repeating sequences of 12 body postures that produced either fluent or non-fluent apparent motion. Importantly, these sequences were symmetrical, with the second half of each sequence mirroring the first half played in reverse (Figure 1). According to the logic of frequency tagging (Norcia et al., 2015), this procedure should result in brain responses at three hierarchical frequencies: a response coupled to individual image presentation, at base rate (BR), a response coupled to the turning point in the sequence, at half cycle rate (BR/6), and a response coupled to the completion of the entire body posture sequence, at full cycle rate (BR/12).

**Figure 1.**
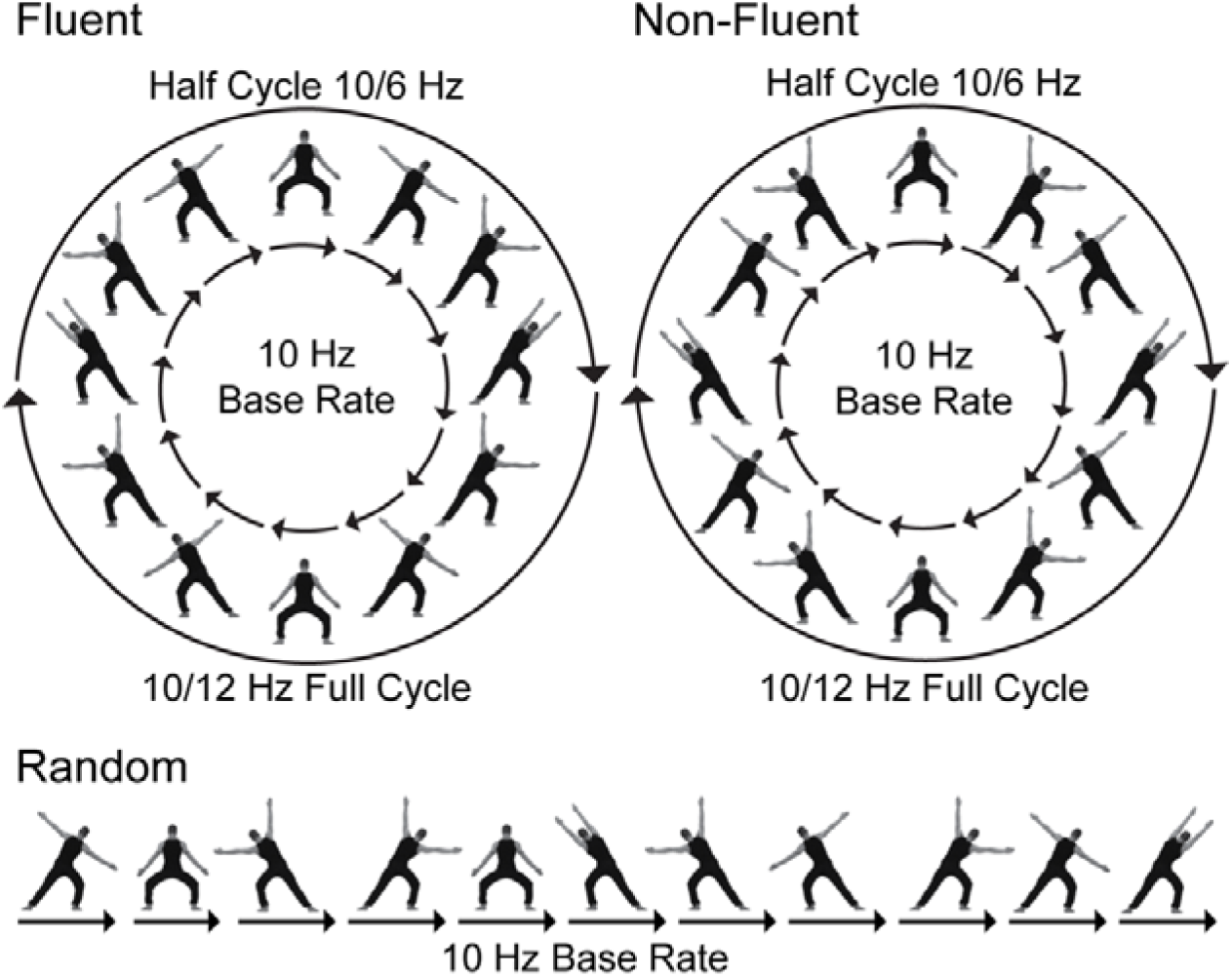
Stimulus sequences for the fluent, non-fluent, and random conditions of Experiment 1. Images are presented at the base rate of 10 Hz. In the fluent condition, images are ordered to induce a coherent movement percept. This percept is perturbed by reordering the images in the non-fluent condition and by showing images at random in the random condition. Fluent and non-fluent sequences have the same symmetrical structure, with the second half of the sequence mirroring the first half in reverse. Hence, in these sequences, a turning point occurs at a frequency of 10/6 Hz (half cycle frequency), and the sequence repeats at a frequency of 10/12 Hz (full cycle frequency).

In Experiment 1 (N = 10), we sought to validate this procedure, and to test if it captures the integration of body postures into movements, for a single agent only (Giese & Poggio, 2003; Lange & Lappe, 2006). To this end, we measured brain activity elicited by fluent, non-fluent, and random sequences (Supplementary Videos; Figure 1). Fluent sequences showed body postures in their natural order. This elicited an apparent motion percept that was perturbed in non-fluent sequences by reordering the postures in a disfluent order and in random sequences by presenting images at random. Hence, all sequences were built from the same postures but differed in their structure. Random sequences did not have any structure and were included only to control for the role of sequential structure and its prediction more generally (Baker et al., 2014). In contrast, fluent and non-fluent sequences both had the same symmetrical structure, but this structure became salient only in fluent sequences. This is because the primary percept in fluent sequences is a series of movements, here presented at half cycle rate, whereas the primary percept in non-fluent sequences is a series of body postures, here presented at full cycle rate (Orgs et al., 2011, 2013, 2016; Orgs & Haggard, 2011; Shiffrar & Freyd, 1990). As a result, if our task captures the temporal binding of bodies into movements, fluent sequences should drive brain activity mainly at half cycle frequencies, whereas non-fluent sequences should instead drive brain activity mainly at full cycle frequencies.

In Experiments 2-3, we then extended our paradigm from individual to group movements, by showing not one but four actors, moving either in or out of synchrony. Synchrony is known to signal the size of a group of stimuli (Braddick et al., 2001; Wagemans et al., 2012), purely based on the sustained temporal coupling of their movement trajectories (Brick & Boker, 2011). If the brain recruits the principles of perceptual organization for social perception and binds temporally related movements into holistic group percepts, then movement-related responses should be stronger for synchronous movements, where a clearly defined relationship between the movements allows them to be integrated more easily. Furthermore, if it is indeed movements that are integrated and not just low-level features, this effect should depend on manipulations known to perturb biological movement processing. Therefore, Experiments 2 (N = 19) and 3 (N = 19) explored the degree to which the influence of synchrony depended on seeing feasible movement trajectories (Experiment 2), by comparing fluent and non-fluent sequences (Lange & Lappe, 2007), and on seeing canonical body orientations (Experiment 3), by comparing upright and inverted agents (Troje & Westhoff, 2006).

Finally, by manipulating both image order and body orientation, we could also investigate whether it is the static, *spatial* (upright or inverted) or the dynamic, *temporal* (fluent or non-fluent) features of the stimulus that are linked together. Given that apparent motion perception is a hierarchical process in which movement processing builds on body processing (Giese & Poggio, 2003; Lange & Lappe, 2006, 2007), group binding can operate either on bodies, on movements, or on both. As body inversion perturbs configural body processing (Lange & Lappe, 2007; Reed et al., 2003), it should interfere primarily with the binding of bodies. In contrast, disfluent movement perturbs global motion processing (Downing et al., 2006; Lange & Lappe, 2007) and should interfere with the binding of movements instead.

## Results

### Dissociating Image, Body, and Movement Processing

To test the hypothesis that base rate, half cycle and full cycle responses capture different components of biological motion processing, we submitted the neural responses in Experiment 1 to a region (left posterior, middle posterior, right posterior, or middle central) x condition (random, non-fluent, fluent) repeated measures ANOVA (Figure 2).

**Figure 2:**
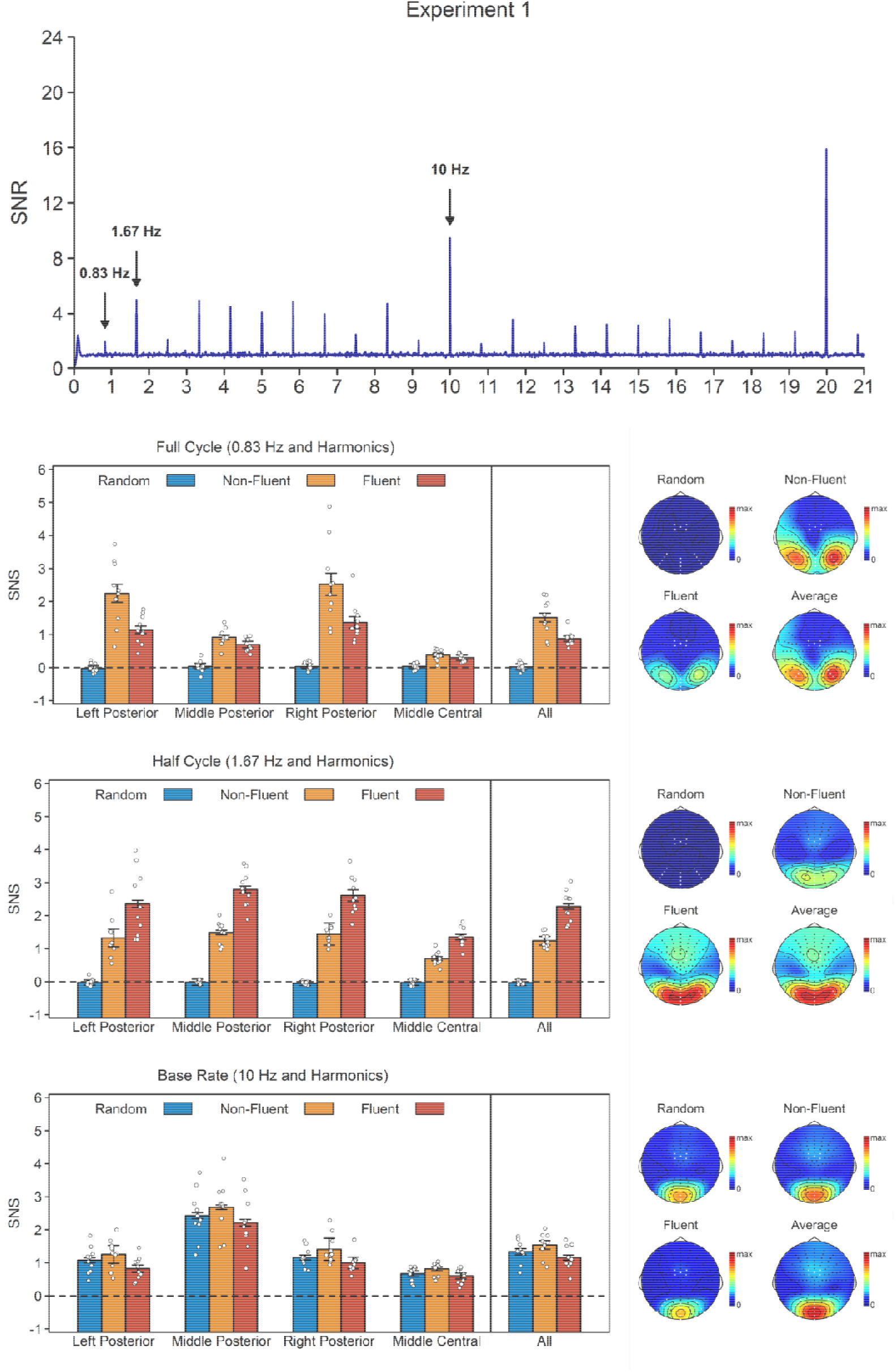
Results of Experiment 1. **Top**. Signal-to-noise ratio (SNR) across all conditions and electrodes of interest. **Bottom**. Noise-subtracted amplitudes (SNS) per condition and response, together with their topographies and an average topography across the three conditions. Topographies are scaled from 0 to the maximum amplitude across the three conditions. The white dots indicate the electrodes included in the analyses. Error bars are standard errors of the mean (SEMs) corrected for within-subject designs (Morey, 2008).

Base rate is coupled to image presentation. As a result, base rate responses should primarily capture the processing of those images. If this is the case, they should be strongest in the non-fluent condition, where the visual change from image to image is largest, and should also be visible in the random condition, where neither body postures nor movements repeat. In line with this hypothesis, the base rate ANOVA revealed a main effect of condition, *F*(2, 18) = 25.23, *p* < .001, η_p_^2^ = .74, with stronger responses in the non-fluent condition than in the random condition, *t*(9) = 4.26, *p* = .002, BF_10_ = 23.67, *d*_z_ = 1.35, and in the random condition than in the fluent condition, *t*(9) = 4.61, *p* = .001, BF_10_ = 36.99, *d*_z_ = 1.46. The base rate response was thus strongest in the non-fluent condition but was also clearly visible in the random condition, *t*(9) = 12.63, *p*_*one-tailed*_ < .001, BF_10_ = 1.30×10^5^, *d*_z_ = 4.00. The focal middle posterior topography further suggested that this response captured relatively low-level visual processing, as demonstrated by a main effect of region, *F*(1.58, 14.25) = 42.57, *p* < .001, η_p_^2^ = .83, indicating that base rate responses were stronger in the middle posterior cluster than in the two lateral posterior clusters, all *t*(9) ≥ 5.21, *p* < .001, BF_10_ ≥ 76.35, *d*_z_ ≥ 1.65, and stronger in the three posterior clusters than in the central cluster, all *t*(9) ≥ 3.73, *p* ≤ .005, BF_10_ ≥ 11.80, *d*_z_ ≥ 1.18.

Full cycle rate is coupled to the completion of the full sequence and full cycle responses should therefore capture the processing of static body sequences. If true, then these responses should be strongest in the non-fluent condition, where the primary percept is a repeating posture sequence. However, unlike the base rate response, they should not be visible in the random condition, as there is no cyclical repetition of the sequences there. In line with this hypothesis, the full cycle ANOVA revealed a main effect of condition, *F*(2, 18) = 52.36, *p* < .001, η_p_^2^ = .85, with stronger responses in the non-fluent than in the fluent condition, *t*(9) = 3.97, *p* = .003, BF_10_ = 16.21, *d*_z_ = 1.25, and stronger responses in the fluent than in the random condition, all *t*(9) = 9.76, *p* < .001, BF_10_ = 5.57×10^3^, *d*_z_ = 3.09. As predicted, full cycle response in the random condition did not differ from baseline, *t*(9) = 0.57, *p*_*one-tailed*_ = .293, BF_10_ = 0.38, *d*_z_ = 0.18. These data thus support the idea that full cycle responses capture the processing of body posture sequences. Consistent with extra-striate visual processing, the full cycle response was characterized by a lateralized posterior topography, as demonstrated by a main effect of region, *F*(2.03, 18.23) = 24.88, *p* < .001, η_p_^2^ = .73, indicating that responses in the two lateral posterior clusters were stronger than responses in all other clusters, all *t*(9) ≥ 4.36, *p* ≤ .002, BF_10_ ≥ 26.97, *d*_z_ ≥ 1.38, and by a condition x region interaction, *F*(2.60, 23.43) = 20.03, *p* < .001, η_p_^2^ = .69, indicating that there was an effect of movement fluency in the left, *t*(9) = 3.96, *p* = .003, BF_10_ = 16.08, *d*_z_ = 1.25, and right, t(9) = 3.59, p = .006, BF_10_ = 9.86, *d*_z_ = 1.14, posterior clusters, but not in the middle posterior, t(9) = 2.09, p = .066, BF_10_ = 1.26, *d*_z_ = 0.66, or central clusters, t(9) = 1.56, p = .153, BF_10_ = 0.65, *d*_z_ = 0.49.

Finally, half cycle rate is coupled to the turning point in the sequence. Importantly, while this point is present both in fluent and non-fluent sequences, it is apparent only in the fluent condition. That is, in the fluent condition, the primary percept is no longer a sequence of postures occurring at full cycle but rather a sequence of movements occurring at half cycle. As a result, if half cycle responses capture the binding of postures into movements, they should be strongest in the fluent condition. Moreover, they should again be absent in the random condition. Supporting this hypothesis, the half cycle analysis revealed a main effect of condition, *F*(2, 18) = 169.53, *p* < .001, η_p_^2^ = .95, with stronger responses in the fluent than in the non-fluent condition, *t*(9) = 7.51, *p* < .001, BF_10_ = 854.42, *d*_z_ = 2.38, and stronger responses in the non-fluent than in the random condition, *t*(9) = 16.19, *p* < .001, BF_10_ = 2.58×10^5^, *d*_z_ = 5.12. Responses in the random condition did not differ from baseline, *t*(9) = - 1.89, *p*_*one-tailed*_ = .954, BF_10_ = 0.10, *d*_z_ = -0.60. Furthermore, suggestive of sensorimotor processing, the half cycle topography included not only by posterior activation, but also a secondary cluster over frontocentral electrodes. This was supported by a main effect of region, *F*(1.53, 13.77) = 13.82, *p* = .001, η_p_^2^ = .61, and a condition x region interaction, *F*(6, 54) = 11.83, *p* < .001, η_p_^2^ = .57, that indicated a stronger response in the three posterior clusters than in the central cluster, all *t*(9) ≥ 3.46, *p* ≤ .007, BF_10_ ≥ 8.25, *d*_z_ ≥ 1.10, and a greater difference between the fluent and non-fluent condition in the middle, *t*(9) = 5.17, *p* < .001, BF_10_ = 73.25, *d*_z_ = 1.64, and right posterior cluster, *t*(9) = 4.86, *p* < .001, BF_10_ = 51.47, *d*_z_ = 1.54, and to a lesser extent in the left posterior cluster, *t*(9) = 2.11, *p* = .065, BF_10_ = 1.28, *d*_z_ = 0.67, than in the central cluster, even though it was significant in all clusters, all *t*(9) ≥ 5.06, *p* < .001, BF_10_ ≥ 64.21, *d*_z_ ≥ 1.60.

In sum, our results clearly dissociate three levels of biological motion processing: Base rate responses captured image processing, full cycle responses captured body sequence processing, and half cycle responses captured movement processing. This was evident both from the pattern across conditions and from the topographies. With regard to the condition pattern, we found that the brain responded most strongly to the half cycle structure of the stimulus when images were presented in sequences producing a coherent movement percept but to its full cycle structure when they were presented in sequences precluding such percept (Orgs et al., 2011, 2013, 2016; Orgs & Haggard, 2011). With regard to the topographies, we found that, consistent with neuroimaging research on low-level visual (Van Essen & Maunsell, 1983), body (Downing, 2001), and movement processing (Caspers et al., 2010), base rate responses were strongest over middle occipital areas, full cycle responses over lateral occipitotemporal areas, and half cycle responses additionally activated frontocentral areas.

### Binding Individual Movements Into Group Movements

Experiment 1 showed that our paradigm could dissociate static and dynamic components of apparent motion processing, thereby capturing the perceptual binding of successive body postures into a continuous movement percept. However, the key question of this study remains to be addressed: how does the brain distinguish multiple people moving independently from multiple people moving in group? Experiments 2 and 3 test the hypothesis that the brain does this by reconstructing not just individual movements from static postures, but by taking into account also the temporal relationships *between* those movements. That is, only synchronous apparent movement trajectories among four individual actors should result in a single percept of group movement. If true, then half cycle responses, which appear to capture the processing of movement trajectories, should not just be sensitive to movement fluency, but also to whether observed movements temporally align across the different actors.

To test this hypothesis, we analyzed half cycle responses in Experiments 2-3 with a repeated measures ANOVA including region, synchrony, and fluency (Experiment 2) or configuration (Experiment 3) as factors (Figures 3-4). In line with our hypothesis, this revealed stronger responses for synchronous than for asynchronous movements in both Experiment 2, *F*(1, 18) = 33.56, *p* < .001, η_p_^2^ = 0.65, and Experiment 3, *F*(1, 18) = 65.77, *p* < .001, η_p_^2^ = 0.79. Experiment 2 also revealed a synchrony x fluency interaction, *F*(1, 18) = 7.36, *p* = .014, η_p_^2^ = 0.29, indicating a stronger synchrony effect for fluent, *t*(18) = 6.13, *p* < .001, *d*_z_ = 1.41, BF_10_ = 2.5×10^3^, than for non-fluent movements, *t*(18) = 3.58, *p* < .001, *d*_z_ = 0.82, BF_10_ = 19.18. In contrast, Experiment 3 revealed a main effect of configuration, *F*(1, 18) = 6.97, *p* = .017, η_p_^2^ = 0.28, with a stronger responses for upright than for inverted dancers, but no synchrony x configuration interaction, *F*(1, 18) = 1.18, *p* = .291, η_p_^2^ = 0.06. In line with Experiment 1, the half cycle topography was characterized by a posterior cluster spreading out over both middle and lateral posterior electrodes and an additional weaker cluster over frontocentral electrodes. This was confirmed by a main effect of region in both Experiment 2, *F*(3, 54) = 78.56, *p* < .001, η_p_^2^ = 0.81, and Experiment 3, *F*(2.13, 38.42) = 53.69, *p* < .001, η_p_^2^ = 0.75, indicating that half cycle responses were significant in all clusters, all *t*(18) > 9.34, *p*_*one-tailed*_ < .001, *d*_z_ ≥ 2.14, BF_10_ ≥ 1.3×10^6^, but stronger over posterior clusters than over the frontocentral cluster, all *t*(18) ≥ 8.34, *p* < .001, *d*_z_ ≥ 1.91, BF_10_ ≥ 1.1×10^5^. None of the other effects reached significance in either experiment.

**Figure 3:**
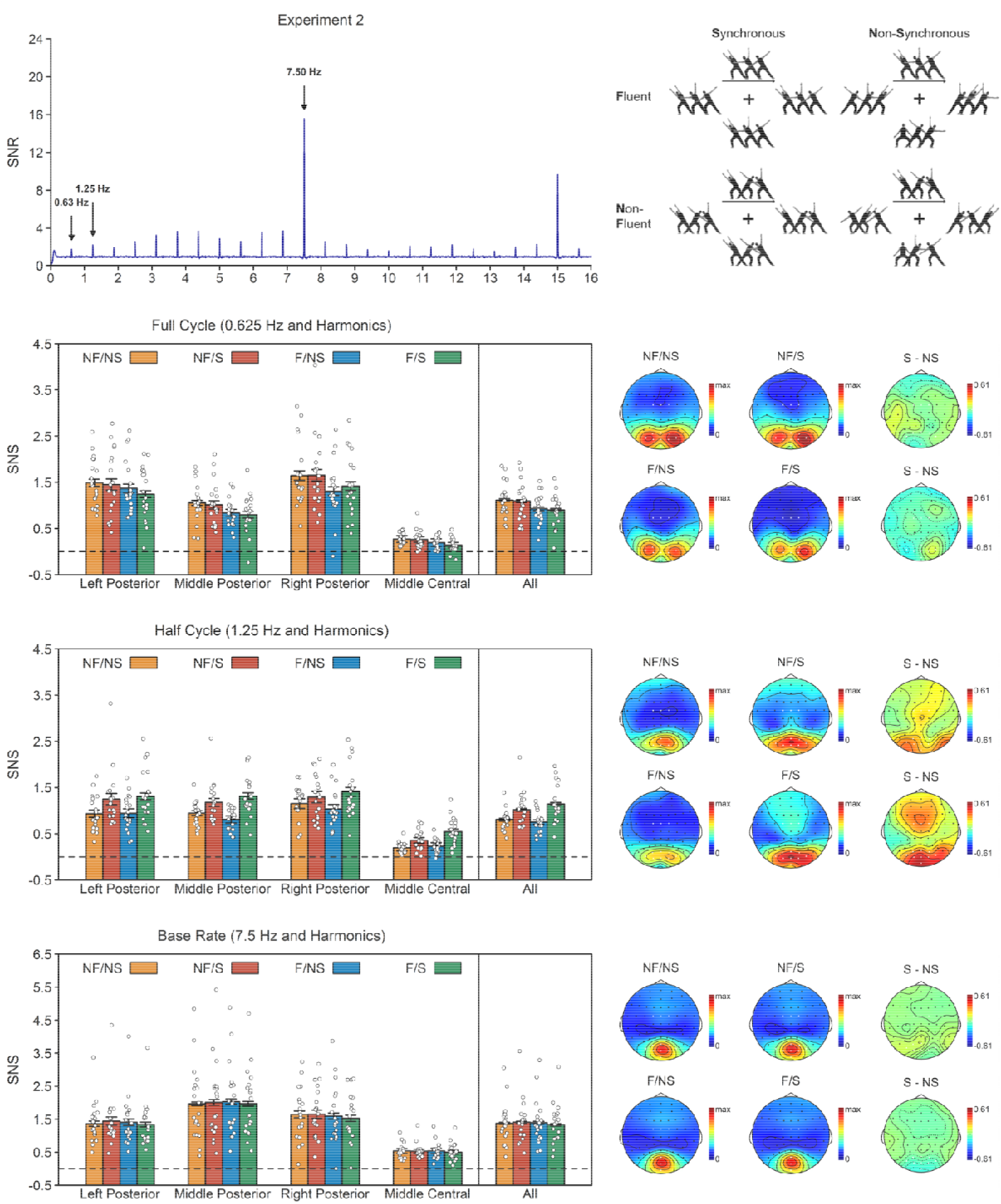
Results of Experiment 2. **Top left**. SNR across conditions and electrodes of interest. Note that Experiments 2 and 3 used a different base rate (7.5 Hz) than Experiment 1 (10 Hz) because this made the asynchronous condition appear more asynchronous. **Top right**. Stimuli of Experiment 2. Participants observed four agents making fluent (F) or non-fluent (NF) movements in synchrony (S) or out of synchrony (NS). **Bottom**. SNS per response and condition, together with their topographies. Topographies are plotted for each of the four conditions and for the synchrony effect on fluent and non-fluent trials. The condition topographies are scaled from 0 to the maximum amplitude across the four conditions. The synchrony effect topographies are scaled symmetrically around zero, with the outer values based on the maximum synchrony effect across topographies. The white dots indicate the electrodes included in the analyses. Error bars are SEMs corrected for within-subject designs (Morey, 2008).

**Figure 4:**
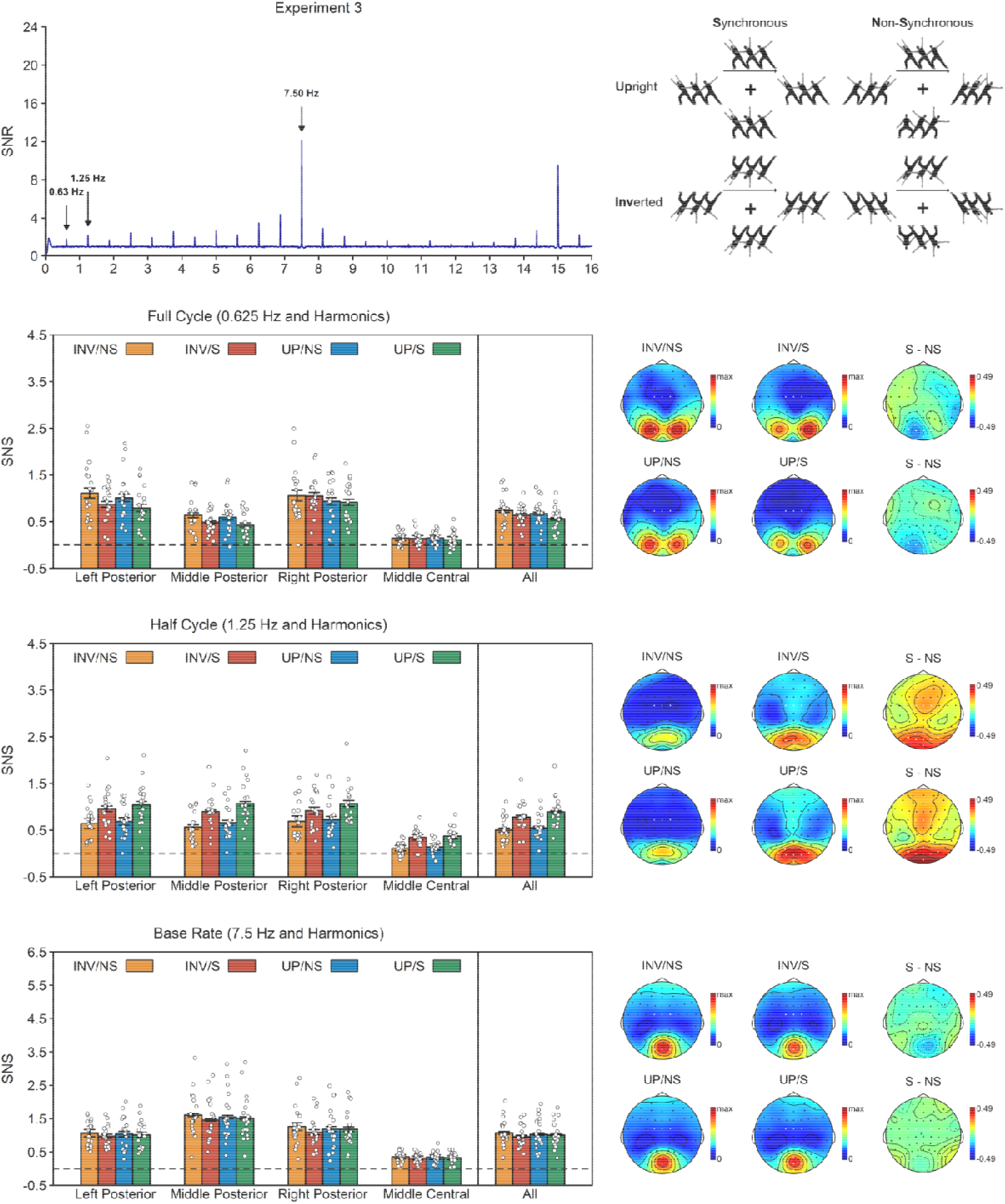
Results of Experiment 3. **Top left.** SNR across conditions and electrodes of interest. Note that Experiments 2 and 3 used a different base rate (7.5 Hz) than Experiment 1 (10 Hz) because this made the asynchronous condition appear more asynchronous. **Top right**. Stimuli of Experiment 3. Participants observed four upright (UP) or inverted (INV) agents moving either in synchrony (S) or out of synchrony (NS). **Bottom**. SNS per response and condition, together with their topographies. Topographies are plotted for each of the four conditions and for the synchrony effect on upright and inverted trials. The condition topographies are scaled from 0 to the maximum amplitude across the four conditions. The synchrony effect topographies are scaled symmetrically around zero, with the outer values based on the maximum synchrony effect across topographies. The white dots indicate the electrodes included in the analyses. Error bars are SEMs corrected for within-subject designs (Morey, 2008).

In contrast to the half cycle results, the full cycle analysis revealed no main effect of synchrony in Experiment 2, *F*(1, 18) = 1.64, *p* = .505, η_p_^2^ = 0.03, and a synchrony effect in the opposite direction in Experiment 3, *F*(1, 18) = 9.29, *p* = .007, η_p_^2^ = 0.34. In addition, Experiment 2 also revealed a main effect of fluency, *F*(1, 18) = 49.37, *p* < .001, η_p_^2^ = 0.73, with stronger responses for non-fluent than for fluent movements, and Experiment 3 revealed a borderline non-significant effect of configuration, *F*(1, 18) = 4.31, *p* = .052, η_p_^2^ = 0.19, with stronger responses for inverted than for upright dancers, but no interactions with synchrony. Similar to Experiment 1, the full cycle topography was characterized by two focal clusters over left and right posterior electrodes, as confirmed by a main effect of region in both Experiment 2, *F*(2.26, 40.66) = 61.53, *p* < .001, η_p_^2^ = 0.77, and Experiment 3, *F*(2.23, 40.14) = 54.90, *p* < .001, η_p_^2^ = 0.75, which indicated that the full cycle response was stronger over left and right posterior clusters than over the other two clusters, all *t*(18) ≥ 4.45, p < .001, *d*_z_ ≥ 1.02, BF_10_ ≥ 102.43. None of the other effects reached significance in either experiment.

Finally, the base rate analysis of Experiment 2 revealed only a main effect of region, *F*(2.06, 37.13) = 30.34, *p* < .001, η_p_^2^ = 0.63. In line with Experiment 1, base rate responses were stronger in the middle posterior cluster than in the other three clusters, all *t*(18) ≥ 2.32, p ≤ .033, *d*_z_ ≥ 0.53, BF_10_ ≥ 1.99. The same region effect was also found in Experiment 3, *F*(2.16, 38.82) = 48.17, *p* < .001. However, Experiment 3 also revealed a reverse synchrony effect, *F*(1, 18) = 7.02, *p* = .016, η_p_^2^ = 0.28, and a region x synchrony x configuration interaction, *F*(3, 54) = 4.28, *p* = .009, η_p_^2^ = 0.19. Post-hoc analyses revealed that this latter effect was driven by a significant synchrony x configuration interaction in the right posterior cluster, *F*(1, 18) = 6.51, *p* = .020, η_p_^2^ = 0.13, but not in the other three clusters, all *F*(1,18) ≤ 3.25, all p ≥ .088, η_p_^2^ ≤ 0.06, indicating stronger responses to asynchronous movements when the actors were inverted, *t*(18) = 3.32, *p* = .004, *d*_z_ = 0.76, but not when they were positioned upright, *t*(18) = 0.04, *p* = .969, *d*_z_ = 0.01. None of the other effects reached significance in either experiment.

## Discussion

This study investigated, in three experiments, the hypothesis that the brain binds temporally related movements into holistic group representations. To this end, we developed a novel approach that combines apparent biological motion with EEG frequency tagging to measure three different components of biological motion processing, early visual image processing (base rate), body sequence processing (full cycle), and movement processing (half cycle), and investigated how these components were modulated by the presence or absence of movement synchrony. Experiment 1 sought to validate our paradigm with individual movements and confirmed that it successfully dissociated between the static body and dynamic movement components of biological motion. Experiments 2 and 3 then extended the paradigm from individual to group movements by showing not one but four actors moving in or out of synchrony in a fluent or non-fluent manner (Experiment 2), in upright or inverted position (Experiment 3).

Half cycle responses in both experiments were stronger for synchronous than for asynchronous movements. In contrast, full cycle and base rate responses were either not sensitive to synchrony (Experiment 2) or stronger for asynchronous movements (Experiment 3). Crucially, the effect of synchrony on half cycle responses was stronger for fluent than disfluent sequences. Given that the internal structure of both sequences was fully matched, this precludes an explanation in terms of low-level processes. Instead, it points towards an interpretation in terms of the temporal relationship between the postures and their apparent movement trajectories. Specifically, our results show that the brain integrates individual movements into group representations by applying the principles of perceptual organization to social perception (Wagemans et al., 2012).

Furthermore, by manipulating body configuration and movement fluency, we could also test specifically which features of biological motion were bound together. That is, body inversion should interfere with the grouping of bodies but not movements (Reed et al., 2003), whereas temporal scrambling should interfere with the grouping of movements but not bodies (Downing et al., 2006). As the modulation of half cycle responses by synchrony was found to depend on movement fluency but not on body inversion, this suggests that it is the dynamic rather than static features of biological motion that are bound into groups. Specifically, our results suggest that synchronous movements are integrated into a single group movement during the temporal integration of body images into movements (Giese & Poggio, 2003; Lange & Lappe, 2006), so that rather than having to analyze the movements of all individual actors, a more efficient movement analysis can take place at the group level. Importantly, however, the half cycle response as such did show an inversion effect. This shows that half cycle responses captured a body-specific process, consistent with the idea that they are the output of a sequential process in which temporal integration of postures into movements occurs only after configural processing of static bodies (Giese & Poggio, 2003; Lange & Lappe, 2006, 2007).

The full cycle response behaved opposite to the half cycle response: whenever the latter was increased, such as when seeing synchronous, fluent, or upright movement, the former was decreased (Supplementary Analysis). This suggests that perturbing movement processing in apparent biological motion causes frame-by-frame processing to take over. The finding that full cycle responses were, if anything, stronger rather than weaker for inverted bodies further indicates that this frame-by-frame processing does not reflect configural body processing (Giese & Poggio, 2003; Lange & Lappe, 2006, 2007; Reed et al., 2003), but rather an earlier, more local analysis of body postures. This interpretation is consistent with the lateral-occipital topography of the full cycle response, pointing towards the extra-striate body area (EBA) as underlying source. Indeed, both brain imaging (Brandman & Yovel, 2010) and brain stimulation (Urgesi et al., 2007) studies have shown that EBA is not sensitive to body inversion. Instead, global body processing has been linked to superior parietal and premotor areas (Urgesi et al., 2007). Similarly, in the current study, the half cycle response, which was perturbed for inverted bodies, also showed a frontocentral component in addition to a posterior component.

Thus, our results show that full cycle responses capture local processing of body posture sequences, whereas half cycle responses capture global processing of body postures, together with the integration of these postures into movements (Giese & Poggio, 2003; Lange & Lappe, 2006). Importantly, this indicates that, even though both responses are harmonically related, they captured distinct processes. This is consistent with evidence that harmonically related components of a musical rhythm such as beat and meter (Nozaradan et al., 2011), beat and rhythmic tapping (Nozaradan et al., 2015), or different meters (Chemin et al., 2014) can likewise produce dissociable responses. Adding onto this, the finding that results were comparable at two different base rates (10 Hz and 7.5 Hz) and for both foveal (Experiment 1) and peripheral (Experiments 2 and 3) stimulus presentation further excludes the possibility that they could be explained purely based on processing basic visual features at a specific presentation frequency.

Finally, the base rate response followed the full cycle response, with either no synchrony effect (Experiment 2) or stronger responses for asynchronous movements (Experiment 3). Interestingly, this finding goes against the results of a recent study that used EEG frequency tagging to measure brain responses to periodic contrast changes of four point-light dancers moving in or out of sync (Alp et al., 2017). The results showed that posterior, occipital brain areas responded more strongly to contrast changes when the dancers moved synchronously, thereby suggesting that low-level visual processing is enhanced rather than reduced when observing synchronous movement. However, this study used two frequencies, with the contrast of one half of the dancers changing at F1 and the contrast of the other half changing at F2. This generates not only fundamental responses at the stimulation frequencies but also intermodulation responses at linear combinations of those frequencies (e.g., F1 + F2), reflecting non-linear neural interactions between the two input streams (Norcia et al., 2015; Zemon & Ratcliff, 1984). In Alp et al. (2017), synchrony modulated the intermodulation but not the stimulation frequencies, indicating that synchrony influenced not early visual processing per se, but rather the integration of early visual features across stimuli. These differences in experimental design can therefore explain why we found reduced rather than increased early visual processing in the asynchronous condition.

By investigating how synchrony drives action perception, the current study has important implications for understanding group alignment (Raafat et al., 2009; Shamay-Tsoory et al., 2019) and its social consequences (Rennung & Göritz, 2016). In particular, our results indicate that multiple people acting together as a group form a strong visual trigger to which the brain entrains more easily than to multiple people acting individually. Furthermore, while speculative, the frontocentral activation cluster observed for half cycle responses could indicate that this neural entrainment may have not only a visual but also a sensorimotor component. Regardless of motor involvement, however, increased neural alignment provides a neurologically feasible explanation for why both human (Dyer et al., 2009; Raafat et al., 2009) and non-human animals (Couzin, 2018; Sumpter, 2006) tend to move in line with the group.

At the same time, increased neural entrainment may also help explain why synchrony is aesthetically pleasing (Eskenazi et al., 2015; Vicary et al., 2017) and a signal of group cohesion (Lakens & Stel, 2011; Marques-Quinteiro et al., 2019; Wilson & Gos, 2019), as stimuli that are processed more fluently are known to produce a hedonic response (Reber et al., 2004). In line with this view, subjective judgements acquired after Experiments 2-3 showed that participants liked the synchronous videos more than the asynchronous videos and Spearman correlations across these two experiments showed that the difference in liking between both conditions correlated positively with the difference in perceived synchrony, ρ = 0.44, *p* = .006, but negatively with the difference in perceived complexity, ρ = -0.42, *p* = .008 (Figure 5).

**Figure 5:**
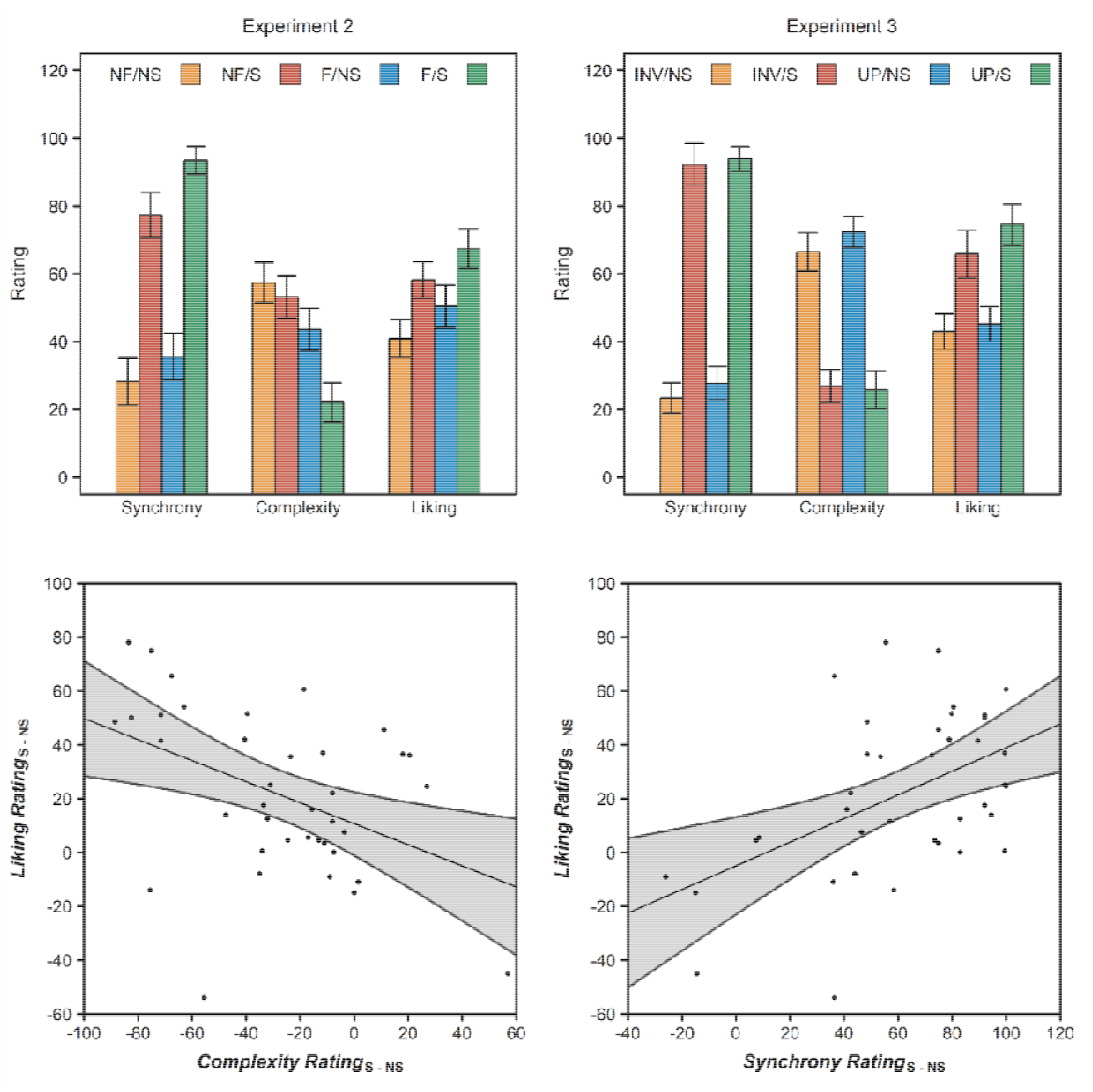
Rating data of Experiments 2 and 3. **Top.** Synchrony, complexity, and liking ratings of fluent (F), non-fluent (NF), upright (UP), inverted (INV), synchronous (S), or non-synchronous (NS) stimuli in Experiments 2 and 3. Error bars are SEMs corrected for within-subject designs (Morey, 2008). **Bottom**. Scatter plots across Experiments 2 and 3 showing how the difference in perceived complexity (left) and perceived synchrony (right) between synchronous and non-synchronous stimuli correlates with the difference in liking between synchronous and non-synchronous stimuli. The shaded grey area shows the 95% confidence interval of a regression.

Finally, our study adds onto a growing body of research investigating ensemble processes in social perception. This research found that two individuals are represented as a single unit when they are either facing (Papeo et al., 2017; Vestner et al., 2019) or interacting (Ding et al., 2017; Liu et al., 2018), and that this changes both how these stimuli are perceived (Liu et al., 2018; Papeo et al., 2017; Vestner et al., 2019) and remembered (Ding et al., 2017; Vestner et al., 2019). By showing that, despite identical input, the brain entrains more strongly to people moving together than to people moving independently, the current results reveal a potential neural mechanism of how ensemble processes shape social perception and evaluation.

To conclude, the current research makes two important contributions. First, it introduces a new approach to study biological motion perception that captures image processing (base rate), local body processing (full cycle), and the temporal integration of configural body snapshots into movements (half cycle) at different frequencies of the EEG spectrum. Second, it extends existing models of biological motion processing by showing that the brain not only binds bodies into movements (Giese & Poggio, 2003; Lange & Lappe, 2006) but also binds movements into groups. These results have important implications for understanding group alignment and its social consequences and provide insight into the neural mechanisms through which observers bind interacting individuals into ensembles shaping perception and memory.

## Methods

### Experiment 1

#### Participants

Ten healthy volunteers with normal or corrected-to-normal vision participated in the experiment (9 female, *M*_age_ = 22.33, SD_age_ = 2.12, range_age_ = 19-26). While this allows us to detect only effects *d*_z_ ≥ 0.72 (Lakens et al., 2018), such effects are to be expected considering the high signal-to-noise ratio of EEG frequency tagging (Norcia et al., 2015; Regan, 1966). All participants signed an informed consent before the experiment and were paid 10 Euros in exchange for their participation. Experiment 1 was conducted at the Université Catholique Louvain and was approved by the local ethics committee.

#### Task, Stimuli, and Procedure

The experiment was programmed in MatLab 2009 (Psychtoolbox). In the experiment, participants saw repeating apparent biological motion sequences consisting of 12 grey-scale body presented on a grey background (Figure 1). There were two experimental conditions (fluent and non-fluent) and one control condition (random). In random sequences, images were presented at random. In contrast, fluent and non-fluent sequences both had a fixed, symmetrical structure in which the second half of the sequence mirrored the first half of the sequence presented in reversed order. In fluent sequences, images were arranged to form a rhythmical dance movement representing a dancer moving from left to right and back from right to left. In the non-fluent condition, these same 12 images were rearranged into a sequence with maximum visual displacement between successive body postures. As a result, even though both sequences were symmetrical, this symmetrical structure was salient only in the fluent sequences (Orgs et al., 2011, 2013, 2016; Orgs & Haggard, 2011).

The three conditions were presented block-wise in randomized order, with 5 blocks per condition. Each block consisted of a 120s video with a 10s fade in and 10s fade out. The fluent and non-fluent videos were created by repeating the corresponding 12-image sequence 100 times and the random videos by presenting a random 1200-image sequence (Video S1-S3). To maintain attention, participants were instructed to fixate on a grey cross in the center of the screen and to press the space bar each time its color changed briefly (200 ms) to red (Rossion et al., 2012).

Importantly, the symmetrical structure of the fluent and non-fluent sequences produces three hierarchical levels of frequency tagging (Figure 1). The first level is driven by the fact that presenting images at 10 Hz produces a 10 Hz base rate response in the EEG signal primarily representing the processing of image onset. In contrast, the second and third levels are driven not by image presentation but by the stimulus structure. The second level reflects the fact that every 6^th^ image (half cycle, at 1.67 Hz) signals a turning point in the sequence and the third level that every 12^th^ image (full cycle, at 0.83 Hz) signals the repetition of the complete sequence. Previous research on apparent biological motion has shown that fluent sequences are primarily seen as movements, whereas non-fluent sequences are primarily seen as a series of body postures (Orgs et al., 2011, 2013, 2016; Orgs & Haggard, 2011; Shiffrar & Freyd, 1990). In our task, movements repeat at half cycle, whereas body posture sequences repeat at full cycle. Therefore, if our task measures the integration of body postures into movements (Giese & Poggio, 2003), fluent sequences should drive brain activity at half cycle frequencies and non-fluent sequences at full cycle frequencies. Moreover, in the random condition, where neither posture sequences nor movements repeat, both half- and full cycle responses should be absent.

#### EEG recording and preprocessing

EEG was recorded from 128 Ag/AgCl active electrodes using a Biosemi EEG system and a sampling rate of 512 Hz. Vertical and horizontal eye movement were measured using four additional electrodes placed on the outer canthus of each eye and in the inferior and superior areas of the right orbit. During EEG recording, all electrodes were referenced to AFz, and electrode impedances were kept below 10 kΩ. All EEG data was offline processed using Letswave 6 (https://www.letswave.org/). Raw data was offline band-pass filtered using a fourth-order Butterworth filter with cut-off values of 0.1 – 100 Hz and segmented according to the experimental conditions (−2 to 122 s). Next, eye movement artefacts were removed by applying ICA on the merged segmented data. Specifically, we analyzed the first 10 components and removed one component for blinks and one or two components related to eye movements. After ICA, faulty or excessively noisy electrodes (< 1% on average) were interpolated using data from the three closest neighboring electrodes. The signal was then re-referenced with respect to the average of all electrodes, before cropping the segments into 96 s epochs (12 to 108 s). At 10 Hz, this ensures that all relevant harmonics are multiples of the epoch duration. Finally, the trials within each condition were averaged and a Fast Fourier Transform (FFT) was applied to transform the data of each electrode to normalized (divided by N/2) amplitudes (µV) in the frequency domain (from 0 to 256 Hz).

#### EEG analysis

For each of the three expected neural responses, at base rate (10 Hz), full cycle (0.83 Hz), and half cycle (1.67 Hz), we extracted the first 10 significant harmonics up until 100 Hz. More specifically, we pooled the electrodes of the grand-averaged signal, computed z-scores for each frequency bin using the 20 surrounding bins except adjacent bins as a baseline, and then selected the first 10 harmonics with z > 2.32 (i.e., *p*_one-tailed_ < 0.01; Retter & Rossion, 2016). Importantly, the three frequencies are harmonically related. Therefore, to minimize overlap, the full cycle response was calculated using only those harmonics that did not overlap with the half cycle harmonics (i.e., the odd harmonics) and the half cycle response using only those harmonics that did not overlap with the base rate harmonics. Accordingly, the full cycle response was calculated as the sum of the amplitudes at 0.83 Hz, 2.50 Hz, 4.17 Hz, 5.83 Hz, 7.50 Hz, 9.17 Hz, 10.83 Hz, 12.50 Hz, 14.17 Hz, and 15.83 Hz. The half cycle response was calculated as the sum of the amplitudes at 1.67 Hz, 3.33 Hz, 5.00 Hz, 6.67 Hz, 8.33 Hz, 11.67 Hz, 13.33 Hz, 15.00 Hz, 16.67 Hz, and 18.33 Hz. Finally, the base rate responses was calculated as the sum of 8 instead of 10 harmonics, 10 Hz, 20 Hz, 30 Hz, 40 Hz, 60 Hz, 70 Hz, 80 Hz, and 90 Hz, excluding electrical noise at 50 Hz and 100 Hz. Importantly, because baseline-subtracted amplitudes were used, the summed response in the absence of signal is expected to be 0 (Retter & Rossion, 2016).

To prevent selection bias, the electrodes entered into the analysis were chosen by averaging the topographies of each response across participants and conditions (Luck & Gaspelin, 2017). This revealed four clusters: a middle posterior cluster with a maximum at Oz, two lateral posterior clusters with maxima at PO7 and PO8, and a frontocentral cluster with a maximum at FCz (Figure 2). The response in each cluster was quantified by taking 5 electrodes centered around the maximum electrode.

The resulting data for each response was analyzed with a condition (fluent, non-fluent, or random) x region (left posterior, middle posterior, right posterior, or middle central) repeated measures ANOVA. ANOVA degrees of freedom were corrected for violation of sphericity using the Greenhouse-Geisser correction whenever Mauchly’s sphericity test was significant (p < .05). Unless otherwise specified, all *t* tests are two-tailed. To further quantify the evidence, all *t* tests are accompanied by Bayes Factors (BFs), calculated with a noninformative Jeffreys prior on the variance and a Cauchy prior with 0 as center and 1 as scale (Rouder et al., 2012). We chose a scale of 1 (“wide prior”) because we expected large effect sizes.

### Experiments 2-3

#### Participants

Two different samples of 20 healthy volunteers participated in Experiment 2 (16 female, *M*_age_ = 26.85, SD_age_ = 6.47, range_age_ = 20-50) and Experiment 3 (14 female, *M*_age_ = 25.80, SD_age_ = 7.10, range_age_ = 18-49). Sample sizes were determined by an a-priori power analysis indicating that 19 participants were necessary to obtain 90% power to detect effect sizes one-third the size of the half cycle fluency effect in Experiment 1. In both experiments, 1 participant had to be excluded because large artefacts across the scalp and throughout the entire experiment made the data uninterpretable. Therefore, the final sample in both experiments comprised 19 participants. All participants signed an informed consent prior to the experiment and were paid £20 in exchange for their participation. Both experiments were conducted at Goldsmiths College, University of London and were approved by the local ethics committee.

#### Task, Stimuli, and Procedure

Both experiments were programmed in PsychoPy (Peirce et al., 2019). The overall procedure was similar to Experiment 1, except that participants now saw not one but four agents, organized in a square grid around the fixation cross, performing fluent or non-fluent movements (Experiment 2) in upright or inverted position (Experiment 3) either in or out of synchrony (Figure 1). Movement fluency was manipulated in the same way as Experiment 1 and synchrony was manipulated by making the agents start from the same (i.e., synchrony) or from different (i.e., asynchrony) starting positions in the sequence (Wilson & Gos, 2019). The 4 starting positions in the asynchronous condition were chosen to maximize perceived asynchrony and were the same for all participants. However, which agent started at which of these 4 positions was counterbalanced across participants.

Experiments 2-3 also used a different presentation rate than Experiment 1. That is, instead of presenting images at a rate of 10 Hz, we used a presentation rate of 7.5 Hz. This was done for two reasons. First, because we wanted to test the degree to which the results of Experiment 1 could be generalized to different frequencies and, second, because a slower presentation rate made the asynchronous condition appear less synchronous. In line with Experiment 1, all conditions were presented block-wise in randomized order, with 5 blocks per condition. However, in contrast to Experiment 1, blocks consisted not of a 120s video but of a 128s video with an 8s fade in and a 8s fade out period. Videos were presented on a white background and were created by repeating the relevant 12-image sequence 80 times (Video S5-S6). To maintain attention and minimize eye movements, participants were asked to focus on a black fixation cross in the center of the screen and to press the space bar each time its color changed briefly (267 ms) to red (Rossion et al., 2012). Before the experiment proper, participants completed 1 practice block where the body postures of all four agents were presented at random, similar to the random condition of Experiment 1. Finally, after the experiment, participants did a brief rating task where they saw a shortened (25s) video of each condition and were asked to rate the synchrony and complexity of the video as well as how much they liked it, on a scale ranging from 0 to 100.

#### EEG recording and preprocessing

EEG was recorded from 64 Ag/AgCl active electrodes using a Biosemi EEG system and a sampling rate of 512 Hz. Vertical and horizontal eye movement were measured using four additional electrodes placed on the outer canthus of each eye and in the inferior and superior areas of the left orbit. During EEG recording, all electrodes were referenced to two electrodes placed on the left and right ear lobes and electrode impedances were kept below 10 kΩ. All EEG data was offline processed using Letswave 6 (https://www.letswave.org/). Raw data was offline band-pass filtered using a fourth-order Butterworth filter with cut-off values of 0.1 – 100 Hz and segmented according to the experimental conditions (−2 to 130s). Next, eye movement artefacts were removed by applying ICA on the merged segmented data, using the same approach as in Experiment 1. After ICA, faulty or excessively noisy electrodes (< 1% on average) were interpolated using data from the three closest neighboring electrodes. In addition, in Experiment 2, a complete block was discarded for a single participant because a large artefact across the scalp disproportionally biased the signal. The signal was then re-referenced with respect to the average of all electrodes, before cropping the segments into 112s epochs (8 to 120s). At 7.5 Hz, this ensures that all relevant harmonics are multiples of the epoch duration. Finally, the trials within each condition were averaged and a Fast Fourier Transform (FFT) was applied to transform the data of each electrode to normalized (divided by N/2) amplitudes (µV) in the frequency domain (from 0 to 256 Hz).

#### EEG analysis

The data was analyzed in the same way as in Experiment 1. Note that this means that the base rate response was now calculated as the sum of the 10 rather than 8 first harmonics because all base rate harmonics were now < 100 Hz. We analyzed activity in the same four electrode clusters as in Experiment 1. However, since we had only 64 instead of 128 electrodes, we used not 5 but 3 electrodes centered around Oz, PO7, PO8, and FCz (Figures 3-4). The resulting data was analyzed with a synchrony (synchronous or asynchronous) x fluency (fluent or non-fluent) x region (left posterior, middle posterior, right posterior, or middle central) repeated measures ANOVA for Experiment 2 and with a with a synchrony (synchronous or asynchronous) x configuration (upright or inverted) x region (left posterior, middle posterior, right posterior, or middle central) repeated measures ANOVA for Experiment 3. All else was equal to Experiment 1.

## Supporting information

Supplementary Analysis

Example Videos Exp1

Example Videos Exp 2

Example Videos Exp 3

## References

Alp, N., Nikolaev, A. R., Wagemans, J., & Kogo, N. (2017). EEG frequency tagging dissociates between neural processing of motion synchrony and human quality of multiple point-light dancers. Scientific Reports, 7(August 2016), 1–9. https://doi.org/10.1038/srep44012

Baker, R., Dexter, M., Hardwicke, T. E., Goldstone, A., & Kourtzi, Z. (2014). Learning to predict: Exposure to temporal sequences facilitates prediction of future events. Vision Research, 99, 124–133. https://doi.org/10.1016/j.visres.2013.10.017

Braddick, O. J., O’Brien, J. M. D., Wattam-Bell, J., Atkinson, J., Hartley, T., & Turner, R. (2001). Brain Areas Sensitive to Coherent Visual Motion. Perception, 30(1), 61–72. https://doi.org/10.1068/p3048

Brandman, T., & Yovel, G. (2010). The Body Inversion Effect Is Mediated by Face-Selective, Not Body-Selective, Mechanisms. Journal of Neuroscience, 30(31), 10534–10540. https://doi.org/10.1523/JNEUROSCI.0911-10.2010

Brick, T. R., & Boker, S. M. (2011). Correlational Methods for Analysis of Dance Movements. Dance Research, 29, 283–304. https://doi.org/10.3366/drs.2011.0021

Caspers, S., Zilles, K., Laird, A. R., & Eickhoff, S. B. (2010). ALE meta-analysis of action observation and imitation in the human brain. NeuroImage, 50(3), 1148–1167. https://doi.org/10.1016/j.neuroimage.2009.12.112

Chemin, B., Mouraux, A., & Nozaradan, S. (2014). Body Movement Selectively Shapes the Neural Representation of Musical Rhythms. Psychological Science, 25(12), 2147–2159. https://doi.org/10.1177/0956797614551161

Couzin, I. D. (2018). Synchronization: The Key to Effective Communication in Animal Collectives. Trends in Cognitive Sciences, 22(10), 844–846. https://doi.org/10.1016/j.tics.2018.08.001

Cracco, E., & Brass, M. (2018a). Automatic imitation of multiple agents: Simultaneous or random representation? Journal of Experimental Psychology: Human Perception & Performance, 44(5), 729–740. https://doi.org/10.1037/xhp0000489

Cracco, E., & Brass, M. (2018b). Motor Simulation of Multiple Observed Actions. Cognition, 180, 200–205. https://doi.org/10.1016/j.cognition.2018.07.007

Cracco, E., & Brass, M. (2018c). The role of sensorimotor processes in social group contagion. Cognitive Psychology, 103, 23–41.

Cracco, E., & Cooper, R. P. (2019). Automatic imitation of multiple agents: A computational model. Cognitive Psychology, 113, 101224. https://doi.org/10.1016/j.cogpsych.2019.101224

Cracco, E., De Coster, L., Andres, M., & Brass, M. (2015). Motor simulation beyond the dyad: Automatic imitation of multiple actors. Journal of Experimental Psychology: Human Perception and Performance, 41(6), 1488–1501. https://doi.org/10.1037/a0039737

Cracco, E., De Coster, L., Andres, M., & Brass, M. (2016). Mirroring multiple agents: Motor resonance during action observation is modulated by the number of agents. Social Cognitive and Affective Neuroscience, 11(9), 1422–1427. https://doi.org/10.1093/scan/nsw059

Cracco, E., Keysers, C., Clauwaert, A., & Brass, M. (2019). Representing Multiple Observed Actions in the Motor System. Cerebral Cortex, 29(8), 3631–3641. https://doi.org/10.1093/cercor/bhy237

Ding, X., Gao, Z., & Shen, M. (2017). Two Equals One: Two Human Actions During Social Interaction Are Grouped as One Unit in Working Memory. Psychological Science, 28(9), 1311–1320. https://doi.org/10.1177/0956797617707318

Downing, P. E. (2001). A Cortical Area Selective for Visual Processing of the Human Body. Science, 293(5539), 2470–2473. https://doi.org/10.1126/science.1063414

Downing, P. E., Peelen, M. V., & Tew, B. D. (2006). The role of the extrastriate body area in action perception. Social Neuroscience, 1(1), 52–62. https://doi.org/10.1080/17470910600668854

Dyer, J. R. G., Johansson, A., Helbing, D., Couzin, I. D., & Krause, J. (2009). Leadership, consensus decision making and collective behaviour in humans. Philosophical Transactions of the Royal Society B: Biological Sciences, 364(1518), 781–789. https://doi.org/10.1098/rstb.2008.0233

Eskenazi, T., Rueschemeyer, S. A., de Lange, F. P., Knoblich, G., & Sebanz, N. (2015). Neural correlates of observing joint actions with shared intentions. Cortex, 70, 90–100. https://doi.org/10.1016/j.cortex.2015.05.027

Giese, M. A., & Poggio, T. (2003). Neural mechanisms for the recognition of biological movements. Nature Reviews Neuroscience, 4(3), 179–192. https://doi.org/10.1038/nrn1057

Grau, A. (2015). Why people dance – evolution, sociality and dance. Dance, Movement & Spiritualities, 2(3), 233–254. https://doi.org/10.1386/dmas.2.3.233_1

Hagen, E. H., & Bryant, G. A. (2003). Music and Dance As a Coalition Signaling System. Human Nature, 14(1), 21–51. https://doi.org/10.1007/s12110-003-1015-z

Hovaidi-Ardestani, M., Saini, N., Martinez, A. M., & Giese, M. A. (2018). Neural Model for the Visual Recognition of Animacy and Social Interaction. In V. Kůrková, Y. Manolopoulos, B. Hammer, L. Iliadis, & I. Maglogiannis (Eds.), Artificial Neural Networks and Machine Learning – ICANN 2018 (Vol. 11141, pp. 168–177). Springer International Publishing. https://doi.org/10.1007/978-3-030-01424-7_17

Jellema, T., & Perrett, D. I. (2003). Perceptual History Influences Neural Responses to Face and Body Postures. Journal of Cognitive Neuroscience, 15(7), 961–971. https://doi.org/10.1162/089892903770007353

Lakens, D., Scheel, A. M., & Isager, P. M. (2018). Equivalence Testing for Psychological Research: A Tutorial. Advances in Methods and Practices in Psychological Science, 1(2), 259–269.

Lakens, D., & Stel, M. (2011). If They Move in Sync, They Must Feel in Sync: Movement Synchrony Leads to Attributions of Rapport and Entitativity. Social Cognition, 29(1), 1–14. https://doi.org/10.1521/soco.2011.29.1.1

Lange, J., & Lappe, M. (2006). A Model of Biological Motion Perception from Configural Form Cues. Journal of Neuroscience, 26(11), 2894–2906. https://doi.org/10.1523/JNEUROSCI.4915-05.2006

Lange, J., & Lappe, M. (2007). The role of spatial and temporal information in biological motion perception. Advances in Cognitive Psychology, 3(4), 419–428. https://doi.org/10.2478/v10053-008-0006-3

Liu, R., Yuan, X., Chen, K., Jiang, Y., & Zhou, W. (2018). Perception of social interaction compresses subjective duration in an oxytocin-dependent manner. ELife, 7, e32100. https://doi.org/10.7554/eLife.32100

Luck, S. J., & Gaspelin, N. (2017). How to get statistically significant effects in any ERP experiment (and why you shouldn’t). Psychophysiology, 54(1), 146–157. https://doi.org/10.1111/psyp.12639

Marques-Quinteiro, P., Mata, A., Simão, C., Gaspar, R., & Farias, A. R. (2019). Observing Synchrony in Dyads: Effects on Observers’ Expectations and Intentions. Social Psychology, 50(3), 174–184. https://doi.org/10.1027/1864-9335/a000377

Morey, R. D. (2008). Confidence intervals from normalized data: A correction to Cousineau (2005). Tutorials in Quantitative Methods for Psychology, 4(2), 61–64. https://doi.org/10.3758/s13414-012-0291-2

Norcia, A. M., Appelbaum, L. G., Ales, J. M., Cottereau, B. R., & Rossion, B. (2015). The steady-state visual evoked potential in vision research: A review. Journal of Vision, 15(6), 4–4. https://doi.org/10.1167/15.6.4.doi

Nozaradan, S., Peretz, I., Missal, M., & Mouraux, A. (2011). Tagging the Neuronal Entrainment to Beat and Meter. Journal of Neuroscience, 31(28), 10234–10240. https://doi.org/10.1523/JNEUROSCI.0411-11.2011

Nozaradan, S., Zerouali, Y., Peretz, I., & Mouraux, A. (2015). Capturing with EEG the Neural Entrainment and Coupling Underlying Sensorimotor Synchronization to the Beat. Cerebral Cortex, 25(3), 736–747. https://doi.org/10.1093/cercor/bht261

Orgs, G., Bestmann, S., Schuur, F., & Haggard, P. (2011). From Body Form to Biological Motion: The Apparent Velocity of Human Movement Biases Subjective Time. Psychological Science, 22(6), 712–717. https://doi.org/10.1177/0956797611406446

Orgs, G., Dovern, A., Hagura, N., Haggard, P., Fink, G. R., & Weiss, P. H. (2016). Constructing Visual Perception of Body Movement with the Motor Cortex. Cerebral Cortex, 26(1), 440–449. https://doi.org/10.1093/cercor/bhv262

Orgs, G., & Haggard, P. (2011). Temporal binding during apparent movement of the human body. Visual Cognition, 19(7), 833–845. https://doi.org/10.1080/13506285.2011.598481

Orgs, G., Kirsch, L., & Haggard, P. (2013). Time perception during apparent biological motion reflects subjective speed of movement, not objective rate of visual stimulation. Experimental Brain Research, 227(2), 223–229. https://doi.org/10.1007/s00221-013-3502-8

Papeo, L., Stein, T., & Soto-Faraco, S. (2017). The Two-Body Inversion Effect. Psychological Science, 28(3), 369–379. https://doi.org/10.1177/0956797616685769

Peirce, J., Gray, J. R., Simpson, S., MacAskill, M., Höchenberger, R., Sogo, H., Kastman, E., & Lindeløv, J. K. (2019). PsychoPy2: Experiments in behavior made easy. Behavior Research Methods, 51(1), 195–203.

Raafat, R. M., Chater, N., & Frith, C. (2009). Herding in humans. Trends in Cognitive Sciences, 13(10), 420–428. https://doi.org/10.1016/j.tics.2009.08.002

Reber, R., Schwarz, N., & Winkielman, P. (2004). Processing Fluency and Aesthetic Pleasure: Is Beauty in the Perceiver’s Processing Experience? Personality and Social Psychology Review, 8(4), 364–382. https://doi.org/10.1207/s15327957pspr0804_3

Reed, C. L., Stone, V. E., Bozova, S., & Tanaka, J. (2003). The Body-Inversion Effect. PSYCHOLOGICAL SCIENCE, 14(4), 7.

Regan, D. (1966). Some characteristics of average steady-state and transient responses evoked by modulated light. Electroencephalography and Clinical Neurophysiology, 20(3), 238–248. https://doi.org/10.1016/0013-4694(66)90088-5

Rennung, M., & Göritz, A. S. (2016). Prosocial Consequences of Interpersonal Synchrony: A Meta-Analysis. Zeitschrift Für Psychologie, 224(3), 168–189. https://doi.org/10.1027/2151-2604/a000252

Retter, T. L., & Rossion, B. (2016). Uncovering the neural magnitude and spatio-temporal dynamics of natural image categorization in a fast visual stream. Neuropsychologia, 91, 9–28. https://doi.org/10.1016/j.neuropsychologia.2016.07.028

Riddell, H., & Lappe, M. (2018). Heading Through a Crowd. Psychological Science, 29(9), 1504–1514. https://doi.org/10.1177/0956797618778498

Rossion, B., Prieto, E. A., Boremanse, A., Kuefner, D., & Belle, G. V. (2012). A steady-state visual evoked potential approach to individual face perception: Effect of inversion, contrast-reversal and temporal dynamics. Neuroimage, 63, 1585–1600.

Rouder, J. N., Morey, R. D., Speckman, P. L., & Province, J. M. (2012). Default Bayes factors for ANOVA designs. Journal of Mathematical Psychology, 56(5), 356–374. https://doi.org/10.1016/j.jmp.2012.08.001

Shamay-Tsoory, S. G., Saporta, N., Marton-Alper, I. Z., & Gvirts, H. Z. (2019). Herding Brains: A Core Neural Mechanism for Social Alignment. Trends in Cognitive Sciences, 23(3), 174–186. https://doi.org/10.1016/j.tics.2019.01.002

Shiffrar, M., & Freyd, J. J. (1990). Apparent Motion of the Human Body. Psychological Science, 1(4), 257–264. https://doi.org/10.1111/j.1467-9280.1990.tb00210.x

Singer, J. M., & Sheinberg, D. L. (2010). Temporal Cortex Neurons Encode Articulated Actions as Slow Sequences of Integrated Poses. Journal of Neuroscience, 30(8), 3133–3145. https://doi.org/10.1523/JNEUROSCI.3211-09.2010

Stevens, J. A., Fonlupt, P., Shiffrar, M., & Decety, J. (2006). New aspects of motion perception: Selective neural encoding of apparent human movements. NeuroReport, 11(1), 109–115.

Sumpter, D. J. (2006). The principles of collective animal behaviour. Philosophical Transactions of the Royal Society of London. Series B, Biological Sciences, 361(1465), 5–22. https://doi.org/10.1098/rstb.2005.1733

Sweeny, T. D., Haroz, S., & Whitney, D. (2013). Perceiving group behavior: Sensitive ensemble coding mechanisms for biological motion of human crowds. Journal of Experimental Psychology: Human Perception and Performance, 39(2), 329–337. https://doi.org/10.1037/a0028712

Tarr, B., Launay, J., Cohen, E. E. A., & Dunbar, R. (2015). Synchrony and exertion during dance independently raise pain threshold and encourage social bonding. Biology Letters, 11, 20150767–20150767. https://doi.org/10.1098/rsbl.2015.0767

Templeton, A., Drury, J., & Philippides, A. (2018). Walking together: Behavioural signatures of psychological crowds. Royal Society Open Science, 5(7), 180172. https://doi.org/10.1098/rsos.180172

Troje, N. F., & Westhoff, C. (2006). The Inversion Effect in Biological Motion Perception: Evidence for a “Life Detector”? Current Biology, 16(8), 821–824. https://doi.org/10.1016/j.cub.2006.03.022

Tsai, J. C. C., Sebanz, N., & Knoblich, G. G. (2011). The GROOP effect: Groups mimic group actions. Cognition, 118(1), 135–140. https://doi.org/10.1016/j.cognition.2010.10.007

Urgesi, C., Calvo-Merino, B., Haggard, P., & Aglioti, S. M. (2007). Transcranial Magnetic Stimulation Reveals Two Cortical Pathways for Visual Body Processing. Journal of Neuroscience, 27(30), 8023–8030. https://doi.org/10.1523/JNEUROSCI.0789-07.2007

Van Essen, D. C., & Maunsell, J. H. R. (1983). Hierarchical organization and functional streams in the visual cortex. Trends in Neurosciences, 6, 370–375. https://doi.org/10.1016/0166-2236(83)90167-4

Vestner, T., Tipper, S. P., Hartley, T., Over, H., & Rueschemeyer, S.-A. (2019). Bound Together: Social Binding Leads to Faster Processing, Spatial Distortion, and Enhanced Memory of Interacting Partners. Journal of Experimental Psychology: General, 148(7), 18.

Vicary, S., Sperling, M., Zimmermann, J. V., Richardson, D. C., & Orgs, G. (2017). Joint action aesthetics. 1–21. https://doi.org/10.1371/journal.pone.0180101

von Zimmermann, J., Vicary, S., Sperling, M., Orgs, G., & Richardson, D. C. (2018). The Choreography of Group Affiliation. Topics in Cognitive Science, 10(1), 80–94. https://doi.org/10.1111/tops.12320

Wagemans, J., Elder, J. H., Kubovy, M., Palmer, S. E., Peterson, M. A., Singh, M., & von der Heydt, R. (2012). A century of Gestalt psychology in visual perception: I. Perceptual grouping and figure–ground organization. Psychological Bulletin, 138(6), 1172–1217. https://doi.org/10.1037/a0029333

Weimerskirch, H., Martin, J., Clerquin, Y., Alexandre, P., & Jiraskova, S. (2001). Energy saving in flight formation. Nature, 413, 697–698.

Wilson, S., & Gos, C. (2019). Perceiving Social Cohesion: Movement Synchrony and Task Demands Both Matter. Perception, 48(4), 316–329. https://doi.org/10.1177/0301006619837878

Zemon, V., & Ratcliff, F. (1984). Intermodulation components of the visual evoked potential: Responses to lateral and superimposed stimuli. Biological Cybernetics, 50, 401–408.

