## Supplementary Analysis for "EEG frequency tagging reveals neural entrainment to people moving in synchrony"

### Comparing Full and Half Cycle Responses

To test the hypothesis that full and half cycle responses were inversely modulated by our manipulations, we tested how amplitudes at these two responses in the left and right posterior electrode clusters were influenced by movement fluency (Experiments 1 and 2), synchrony (Experuments 2 and 3), and body configuration (Experiment 3). Note that we focused on the left and right posterior clusters because these were the only two clusters that were consistently activated for both responses. The results of a frequency (half cycle vs. full cycle) x fluency (fluent vs. non-fluent) x region (left vs. right posterior) repeated measures ANOVA on the data of **Experiment 1** revealed only a frequency x fluency interaction, $F(1, 9) = 37.05$, $p < .001$, $\eta_p^2 = .80$, indicating that the full cycle response was reduced, $t(9) = -4.13$, $p = .003$, $BF_{10} = 20.14$, $d_z = 1.30$, but the half cycle response increased, $t(9) = 6.85$, $p < .001$, $BF_{10} = 452.66$, $d_z = 2.17$, on fluent compared with non-fluent trials.

A frequency x synchrony (synchrony vs. asynchrony) x fluency x region repeated measures ANOVA on the data of **Experiment 2** likewise revealed a frequency x fluency interaction, $F(1, 18) = 10.42$, $p = .005$, $\eta_p^2 = .37$, with smaller full cycle responses, $t(18) = -6.80$, $p < .001$, $BF_{10} = 9.8x10^3$, $d_z = 1.56$, but not half cycle responses, $t(18) = 0.27$, $p = .787$, $BF_{10} = 0.18$, $d_z = 0.06$, on fluent compared with non-fluent trials, as well as a frequency x synchrony interaction, $F(1, 18) = 21.50$, $p < .001$, $\eta_p^2 = .54$, with larger half cycle responses, t(18) = 4.56, p < .001, BF10 = 132, dz = 1.05, but not full cycle responses, $t(18) = -0.23$, $p = .818$, $BF_{10} = 0.18$, $d_z = 0.05$, on synchronous compared with asynchronous trials.

Finally, a frequency x synchrony x configuration (upright vs. inverted) x region repeated measures ANOVA on the data of **Experiment 3** revealed a frequency x synchrony interaction, $F(1, 18) = 34.46$, $p < .001$, $\eta_p^2 = .66$, with increased half cycle responses, $t(18) = 8.02$, $p < .001$, $BF_{10} = 8.2x10^4$, $d_z = 1.84$, but reduced full cycle responses, $t(18) = -2.34$, $p =$

.031, $BF_{10}$ = 1.76, $d_z$ = 0.54, on synchronous compared with asynchronous trials, and a frequency x configuration interaction, $F(1, 18) = 15.70$, $p = .001$, $\eta_p^2 = .47$, with larger half cycle responses, $t(18) = 2.12$, $p = .049$, $BF_{10}$ = 1.20, $d_z$ = 0.49, but weaker full cycle responses, $t(18) = -2.41$, $p = .027$, $BF_{10}$ = 1.96, $d_z$ = 0.55, on upright compared with inverted trials.

Taken together, these results indicate that manipulations that tended to strengthen the half cycle response also tended to weaken the full cycle response. This is consistent with the hypothesis that these responses reflect dissociable processes, despite being harmonically related.
